# AGPAT1 is a novel Chikungunya virus receptor on human cells

**DOI:** 10.1101/2025.05.20.655106

**Authors:** Brohmomoy Basu, Debapriyo Sarmadhikari, Shailendra Asthana, Manjula Kalia, Sudhanshu Vrati

## Abstract

Chikungunya virus (CHIKV) is a medically important alphavirus whose host receptor is not fully understood. We identified AGPAT1 from the human Huh7 cell plasma membrane binding with CHIKV particles in vitro. The CHIKV binding with AGPAT1 was demonstrated on Huh7, HAP1, and ERMS plasma membrane by confocal microscopy. The AGPAT1 antibody inhibited CHIKV binding to cells, reducing the virus uptake in Huh7 and ERMS cells. CHIKV binding, uptake, and replication were significantly reduced in the AGPAT1 knockout HAP1 cells, and the ectopic expression of AGPAT1 rescued the reduced virus binding, uptake, and replication. AGPAT1 interacted with the E1 surface on the E1-E2 dimer of CHIKV envelope proteins *in silico*. The E1-AGPAT1 interacting amino acid residues identified in the computational study were experimentally verified. AGPAT1 was also shown to have a role in the binding and uptake of another alphavirus, Ross River virus. These data demonstrate an important role for AGPAT1 as a novel host receptor involved in CHIKV binding and uptake in human cells.

**AUTHOR SUMMARY:** Chikungunya virus (CHIKV) is a medically important alphavirus. Identifying its receptor on human cells will aid the development of novel antivirals. We have identified AGPAT1 as a novel CHIKV receptor on several human cells. We provide data showing the interaction of AGPAT1 with the E1 protein of CHIKV E1-E2 dimer present on the virion surface and identify the amino acid residues involved in the interaction. AGPAT1 was also shown to have a role in the binding and uptake of Ross River virus, another alphavirus. AGPAT1 is a protein involved in lipid metabolism and has not been reported to have a direct role in any virus infection, or as a virus receptor. Thus, this work identifies AGPAT1 as a novel receptor for CHIKV on human cells, and has implications for understanding the virus pathogenesis and designing the receptor-blocking CHIKV antivirals.

## INTRODUCTION

Chikungunya virus (CHIKV) is an Alphavirus transmitted mainly by *Aedes aegypti* and *Aedes albopictus* mosquitoes (1). CHIKV causes high fever, maculopapular rashes, debilitating pain, and polyarthralgia. A major outbreak of chikungunya fever occurred in India in 2006, where 1.25 million cases were reported (2). In the last fifteen years, the virus has spread to over 110 countries in Asia, Europe, South and Central America, and Africa (3). Although CHIKV is not a deadly virus, it causes severe, long-lasting indisposition in a large population each year and is thus considered a priority pathogen. A live, attenuated CHIKV vaccine was approved recently by the FDA (4, 5). However, no CHIKV-specific antivirals are available.

CHIKV has a single-stranded, positive-sense, ∼12 kb long RNA genome encoding four non-structural and five structural proteins (C, E3, E2, 6K, and E1). The dimers of the E1 and E2 proteins, ultimately forming trimers, are present as spikes on the virion surface (6). The E1 protein, a 47-kDa transmembrane protein, is involved in the fusion step during infection. The 50-kDa transmembrane E2 protein induces neutralizing antibodies (7, 8). Both E1 and E2 are involved in receptor binding (9–11).

CHIKV infects different animal hosts such as mosquitoes (12), humans (13), monkeys (14), rodents (15), bats (16), birds (17), and livestock animals (18). Besides, the virus can infect a variety of cell types, such as fibroblasts (19), myocytes (20), epithelial cells (19), macrophages (21), and dendritic cells (22). The variety of CHIKV-susceptible host species and cell types suggests that the virus might use diverse cell surface molecules as an attachment factor or receptor.

Several attachment factors have been identified for alphaviruses. These include glycosaminoglycans (GAGs) such as heparan sulfate (23) and chondroitin sulfate (24). The 181/25 CHIKV isolate needed heparan sulfate for its infectivity, whereas the LR2006 OPY1 CHIKV strain showed dependence upon heparan sulfate proteoglycans (HSPGs) or other GAGs for infectivity (22). While the cell-surface GAGs promote viral entry, they are not essential for CHIKV infection. CHIKV can enter GAG-deficient cells, suggesting GAG-independent entry pathways exist (25). Thus, the attachment factor, GAGs, mainly enhances the virus infection.

Studies have identified several potential cell surface receptors facilitating CHIKV entry into the host cell (26). Thus, the Matrix Remodeling Associated 8 (MXRA8) protein serves as a receptor for CHIKV and other arthritogenic alphaviruses in mouse and human cells (27). Another protein, Prohibitin, was shown to be involved in the uptake of CHIKV in the human microglial CHME-5 cells (28). In 293T cells, the CD147 protein complex acted as a possible entry factor for CHIKV and other alphaviruses (29). Besides, TIM-1 and DC-SIGN have been shown as the CHIKV attachment factors on mammalian cells (30, 31).

These studies illustrate the complex interactions between CHIKV and the host cell surface proteins, enabling viral entry and infection. None of the proteins described above serve as the exclusive receptor for CHIKV entry into the host cell, and more CHIKV receptor proteins remain to be discovered. Identification of CHIKV receptors in mammalian cells will help develop the virus entry inhibitors as novel antivirals. Here, we describe the 1-acyl-sn-glycerol-3-phosphate acyltransferase alpha (AGPAT1) protein that acts as a novel CHIKV receptor on Huh7, HAP1, and ERMS human cells.

## RESULTS

### CHIKV binds AGPAT1 protein from the Huh7 plasma membrane *in vitro*

The Huh7 plasma membrane proteins’ binding with purified CHIKV was carried out *in vitro,* and the interacting host proteins were identified by mass spectrometry. From the six independent experiments, the CHIKV-binding proteins identified on two or more occasions are shown in Fig. S1. While Junction plakoglobin (JUP) and Flaggrin (FLG) were seen in three, Annexin 2 (ANXA2) and Hornerin (HRNR) in four, AGPAT1 was recorded in five experiments.

### AGPAT1 interacts with CHIKV on the Huh7 plasma membrane *in cellulo*

AGPAT1 is reported on the mammalian cell endoplasmic reticulum membrane (32, 33). To function as a CHIKV cellular receptor, it must be located on the plasma membrane. Indeed, AGPAT1 was located on the Huh7 plasma membrane, as revealed by immunofluorescence imaging of the non-permeabilized cells (Fig. S2A). Western blotting of the cytoplasmic and membrane fractions (devoid of the ER membranes) confirmed the presence of AGPAT1 on the Huh7 plasma membrane (Fig. S2B).

Immunofluorescence-based confocal microscopy showed colocalization (PCC 0.62) of CHIKV virions and AGPAT1 on the plasma membrane (Fig. 1A). The super-resolution structured illumination microscopy corroborated this finding (Fig 1A, lower panel). This was further confirmed by immunoprecipitation of AGPAT1 from the Huh7 plasma membrane fraction incubated with purified CHIKV (Fig. 1B).

**Figure 1.**
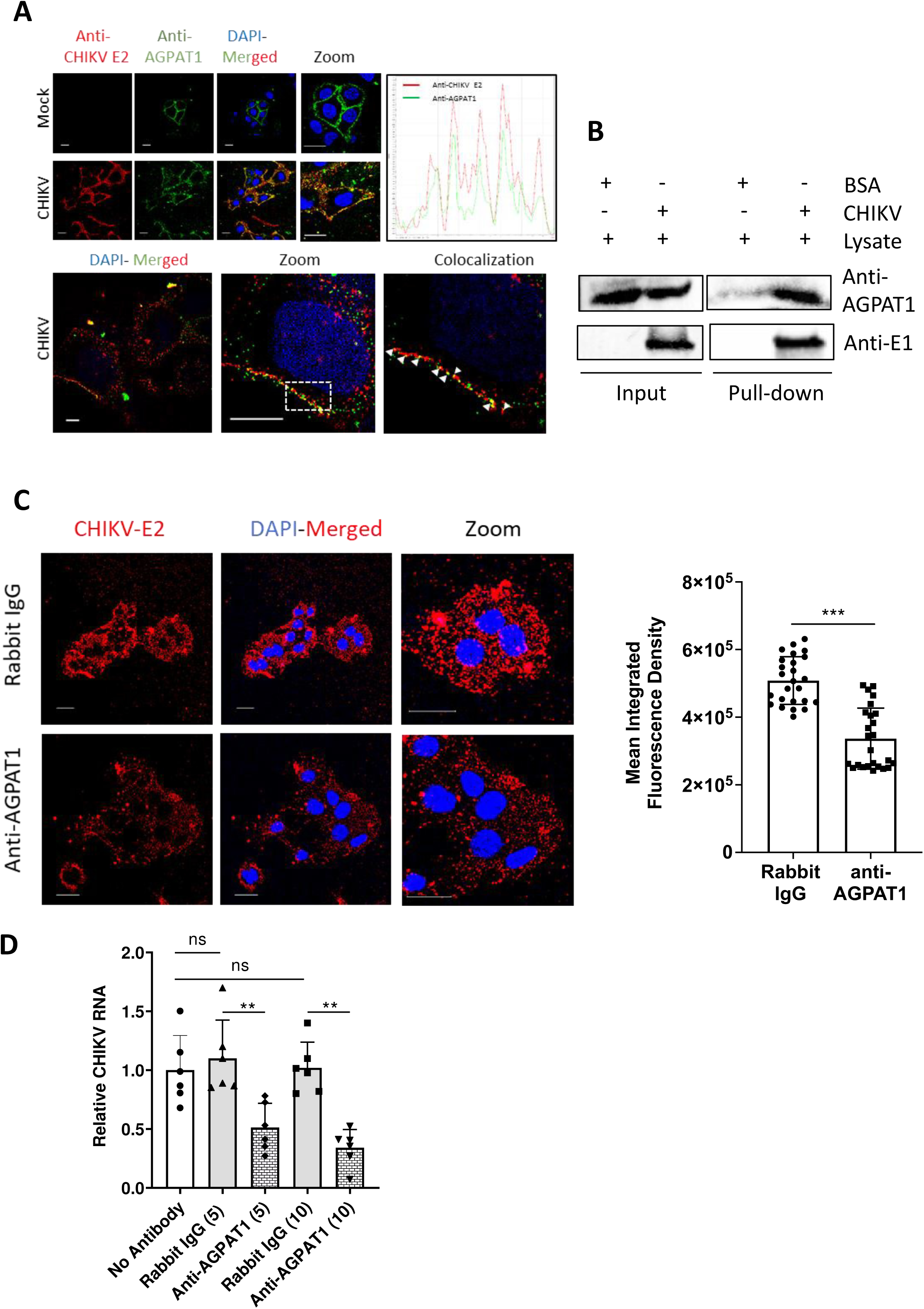
AGPAT1 interacts with CHIKV on the Huh7 plasma membrane. (A) Huh7 cells were incubated with CHIKV (MOI 50) on ice for 45 min, followed by CHIKV anti-E2 mouse monoclonal antibody and anti-AGPAT1 rabbit polyclonal antibody for another 30 min. The cells were washed with ice-cold PBS and incubated with anti-mouse Alexa Fluor-647 and anti-rabbit Alexa Fluor-488 antibodies for 30 min. The nuclei were stained with DAPI, and the cells were imaged under the Leica SP8 confocal microscope, as shown in the top left panel (scale bar = 25 µm). The PCC was determined for the co-localization of Alexa Fluor-488 (AGPAT1) and Alexa Fluor-647 (CHIKV) on the plasma membrane. The top right panel shows the line plot showing the co-localization of the red (CHIKV) and green (AGPAT1) fluorescence signals. The bottom panel shows the colocalization of CHIKV and AGPAT1 observed by the super-resolution structured illumination microscopy (SIM) under the Zeiss Elyra PSI microscope (scale bar = 10 µm). The white arrows show the co-localized puncta under 100X magnification and 10X zoom. The representative images are shown. (B) The Huh7 cell lysate was incubated with the purified CHIKV or BSA (negative control). The immunoprecipitation was carried out using the CHIKV anti-E1 antibody. The input cell lysate and the pull-down were Western blotted with AGPAT1 or CHIKV E1 antibodies. A representative blot is shown. (C) Huh7 cells were incubated on ice with rabbit anti-AGPAT1 polyclonal antibody or rabbit IgG (isotype control) for 30 min at 10 µg/ml concentration, followed by incubation with CHIKV (MOI 50) on ice for 30 min. The cells were then incubated with CHIKV anti-E2 mouse monoclonal antibody on ice for 30 min and washed with ice-cold PBS. The cells were stained with Alexa Fluor-647 anti-mouse antibody for 30 min. The nuclei were stained with DAPI, and the cells were imaged using the Leica SP8 confocal microscope (scale bar = 25 µm). The MIFD is shown as mean±SD in the right panel. The experiment was done in triplicate, and a representative image of the stained cells is shown. The Student’s t-test was conducted to analyze the data; \*\*\**p*<0.001. (D) Huh7 cells were incubated with 5 or 10 mg/ml concentration (indicated in the brackets) of rabbit anti-AGPAT1 antibody or rabbit IgG (isotype control) on ice for 30 min, or without antibody, and then incubated with CHIKV (MOI 1) on ice for 1 h. The cells were then washed with ice-cold PBS and incubated for 3 h at 37 °C. The total RNA from the cells was isolated, and the CHIKV RNA level was determined by qRT-PCR. The CHIKV RNA level in the no-antibody control was taken as 1. The statistical analysis of the data shown as mean±SD was done using the Student’s t-test with Welch’s correction, \*\**p*<0.01.

CHIKV binding to Huh7 cells was studied by measuring the mean integrated fluorescence density (MIFD) on cells incubated with CHIKV, followed by immunofluorescence staining and confocal microscopy. The MIFD was reduced (*p*<0.001) by 34% on cells incubated with CHIKV in the presence of AGPAT1 antibody when compared with the control (Fig. 1C), reinforcing the finding that CHIKV binds to AGPAT1 on Huh7 plasma membrane. Corroborating this, CHIKV infection of Huh7 cells was reduced (*p*<0.01) by 53% and 68% in the presence of 5 µg/ml and 10 µg/ml AGPAT1 antibody in a dose-dependent manner, respectively (Fig. 1D). The rabbit IgG (isotype control) did not affect the virus infection.

### CHIKV uptake in Huh7 cells is related to AGPAT1 protein levels

The siRNA-mediated knockdown of AGPAT1 was used to study its role in CHIKV uptake (Fig. S3A). The siRNA treatment did not affect the cell viability. The CHIKV RNA levels were lowered (*p*<0.001) by 62%, 50%, and 48% in the siAGPAT1-treated CHIKV-infected cells compared to the control at 1, 3, and 6 h pi, respectively, showing a 62% reduced virus uptake (Fig. S3B). The viral titers were 83%, 73%, and 48% lower (*p*<0.001) in the siAGPAT1-treated cells than in the control at 6, 12, and 18 h pi, respectively. Further, CHIKV uptake was studied in Huh7 cells ectopically expressing the HA-tagged protein AGPAT1-HA (Fig. S3C). A significantly enhanced (*p*<0.01) level of viral RNA (175% and 240% higher at 1 and 6 h pi, respectively) was seen in AGPAT1-overexpressing CHIKV-infected cells compared to the empty vector-transfected control cells; showing a 175% enhanced virus uptake. Similarly, higher titers (*p*<0.001) of CHIKV (175%, 240%, and 120% higher at 6, 12, and 18 h pi, respectively) were seen in AGPAT1-overexpressing CHIKV-infected cells (Fig. S3D).

### CHIKV binding and uptake are reduced in the AGPAT1 knockout (KO) HAP1 cells

HAP1 is a near-haploid human cell line. The AGPAT1 KO HAP1 cell line was generated and validated for the absence of AGPAT1 by Western blotting (Fig. 2A). Notably, immunofluorescence staining for AGPAT1 was seen on the plasma membrane of the wild-type (WT) HAP1 cells, while it was absent on the KO cells (Fig. 2B).

**Figure 2.**
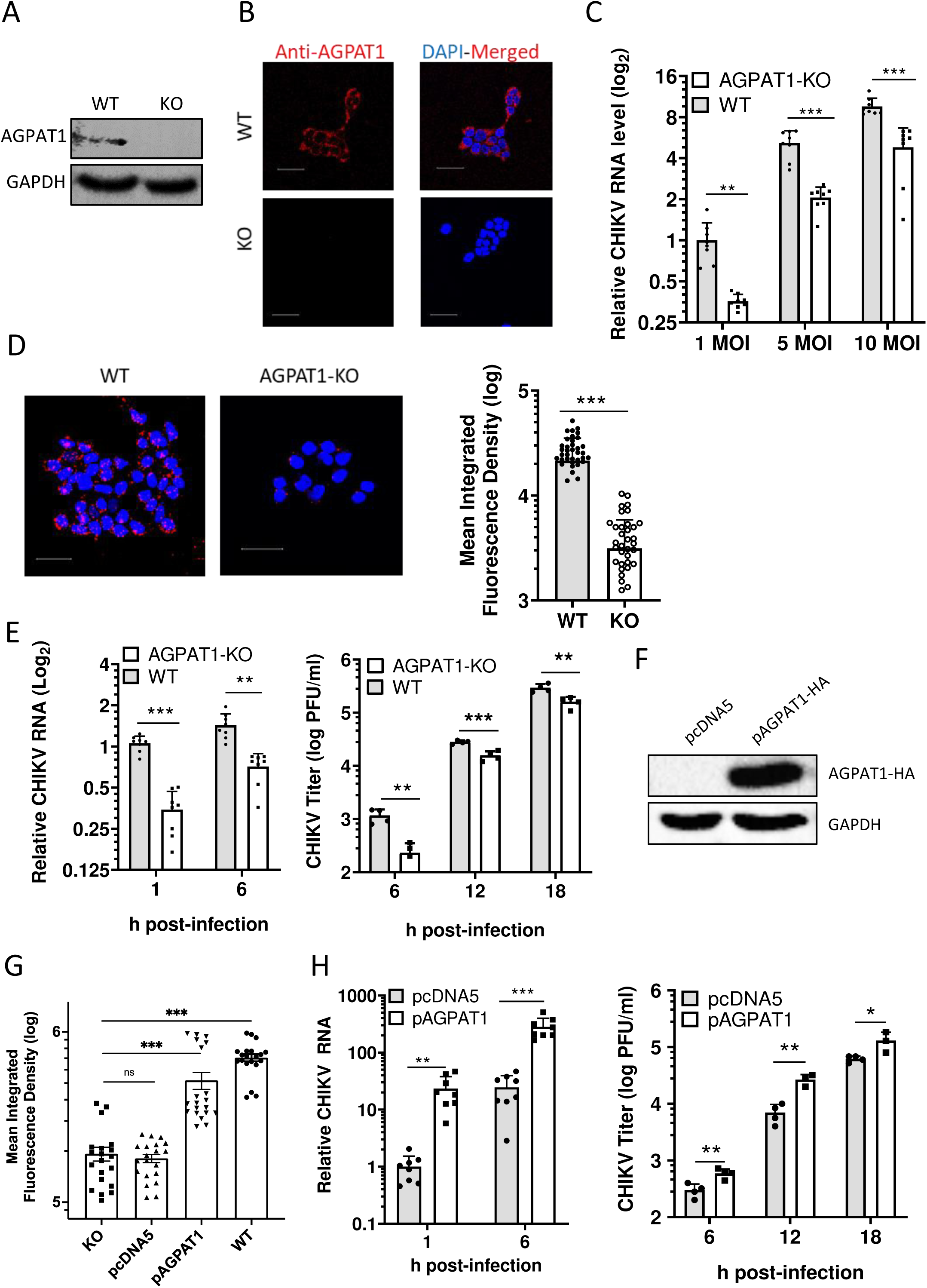
CHIKV binding and uptake in AGPAT1 knockout HAP1 cells. (A) The wild-type (WT) and AGPAT1 knockout (KO) HAP1 cell lysates were Western blotted with anti-AGPAT1 antibody to establish the absence of AGPAT1 in the KO cells. GAPDH was used as the loading control. A representative blot is shown. (B) The HAP1 KO and WT cells were incubated with rabbit anti-AGPAT1 antibody and fixed by 2% paraformaldehyde, followed by incubation with anti-rabbit Alexa Fluor-568 antibody. The nuclei were stained with DAPI, and the cells were imaged using the Leica SP8 confocal microscope (scale bar = 25 µm). The representative images are shown. (C) The KO and WT HAP1 cells were incubated with CHIKV on ice for 30 min at the indicated MOI to allow the virus binding. The unbound virus was removed by ice-cold PBS wash. The cells were then lysed, and total RNA was extracted. The CHIKV binding to cells was determined by the level of CHIKV RNA using qRT-PCR. The relative CHIKV RNA levels are presented where the CHIKV RNA level at 1 MOI in the WT cells was taken as 1. The data from 4 biological replicates and 2 technical replicates are shown. (D) The KO and WT HAP1 cells were incubated with CHIKV (MOI 25) on ice for 30 min. This was followed by incubating the cells with CHIKV anti-E2 mouse antibody for 30 min on ice, and a wash with the ice-cold PBS. The cells were fixed and immunostained with anti-mouse Alexa Fluor-647 antibody. DAPI was used to stain the nuclei. The cells were imaged using the Leica SP8 confocal microscope (scale bar = 25 µm), and the MIFD was determined. The representative images are shown. The MIFD data are from 3 biological replicates. (E) The WT and AGPAT1 KO HAP1 cells were incubated with CHIKV (MOI 0.1) on ice for 1 h. The cells were washed to remove the residual virus and incubated at 37°C. The cells were treated with trypsin to remove the extracellular virion particles before the harvest. The cells and the culture supernatants were harvested at the indicated times. The CHIKV RNA in the cells was quantified by qRT-PCR and the viral titers in the culture supernatant were determined by plaque assay. The relative CHIKV RNA levels are presented (left panel) where the CHIKV RNA level at 1 h pi in the WT cells was taken as 1. The right panel has the virus titers. The data from 4 biological replicates and 2 technical replicates are shown. (F) The AGPAT1-KO HAP1 cells were transfected with pAGPAT1-HA or pcDNA5, and incubated for 48 h at 37 °C. The Western blotting showed the ectopic expression of the AGPAT1-HA protein in the transfected cells. A representative blot is shown. (G) The HAP1 KO cells transfected with pAGPAT1-HA or pcDNA5 as above, were fixed and incubated with CHIKV (MOI 25) on ice for 30 min. This was followed by incubating the cells with CHIKV anti-E2 mouse antibody for 30 min on ice, and a wash with the ice-cold PBS. The cells were immunostained with the anti-mouse Alexa Fluor-647 antibody. DAPI was used to stain the nuclei. The cells were imaged using the Leica SP8 confocal microscope, and the MIFD was determined. The MIFD data from 3 biological replicates are shown. (H) The AGPAT1 KO HAP1 cells were transfected with the plasmid pAGPAT1-HA or pcDNA5 and 48 h later were incubated with CHIKV (MOI 5) for 1 h on ice. The cells were washed with ice-cold PBS and then incubated at 37 °C. The cells and the culture supernatants were harvested at the indicated times. The CHIKV RNA in the cells was quantified by qRT-PCR and the viral titers in the culture supernatant were determined by plaque assay. The relative CHIKV RNA levels are presented (left panel), where the CHIKV RNA level at 1 h pi in the pcDNA5-transfected cells was taken as 1. The right panel has the virus titers. The data from 4 biological replicates and 2 technical replicates are shown. The statistical analysis of the data presented as mean±SD used the Student’s t-test with Welch’s correction; \**p*<0.05, \*\**p*<0.01, \*\*\**p*<0.001.

Compared to the WT HAP1 cells, the KO cells showed 64%, 60%, and 50% reduction (*p*<0.001) in CHIKV binding at MOI 1, 5, and 10, respectively (Fig. 2C). The confocal microscopy (Fig. 2D) showed that CHIKV binding, measured as the MIFD, was ∼85% lower (*p*<0.001) in the KO cells than in the WT cells. CHIKV RNA levels in the KO cells showed 71% reduction both at 1 and 6 h pi compared to the WT cells, showing a 71% reduced virus uptake (Fig. 2E). Notably, the virus replication, measured as the extracellular virus titers, was reduced (*p*<0.001) by 80%, 44%, and 45% at 6, 12, and 18 h pi, respectively, in the KO cells compared to the WT cells (Fig. 2E).

To see if the reduced CHIKV binding, uptake, and replication in the KO HAP1 cells could be rescued by the ectopically-expressed AGPAT1, the cells were transfected with the pAGPAT1-HA expression vector or with the control empty vector pcDNA5. The Western blotting established the expression of AGPAT1-HA in the transfected cells (Fig. 2F). The CHIKV binding, determined by MIFD, was 200% higher (*p*<0.001) in the KO cells ectopically-expressing AGPAT1-HA than in the KO cells (Fig. 2G). A 23- and 12-fold enhanced (*p*<0.01) viral RNA levels were seen in the AGPAT1-expressing KO cells than in the control at 1 and 6 h pi, respectively (Fig. 2H), showing a 2300% enhanced CHIKV uptake. Similarly, CHIKV replication was enhanced (*p*<0.01) in the AGPAT1-expressing HAP1 KO cells. Thus, CHIKV titers were 100%, 300%, and 100% higher in the rescued cells compared to the control at 6, 12, and 18 h pi (Fig. 2H).

These data show that CHIKV binds AGPAT1 on the HAP1 plasma membrane, and the virus binding, uptake, and replication are significantly reduced in the cells lacking AGPAT1.

### *In silico* analysis shows AGPAT1 interaction with CHIKV E1 protein

CHIKV has E1 and E2 proteins on the virion surface that could interact with AGPAT1, together or individually. To obtain the most likely complex, the interaction of the CHIKV E1 and E2 dimer with AGPAT1 was studied *in silico*. A total of 100 protein-protein docking poses were generated using HDOCK. To rule out any HDOCK bias, complexes using AlphaFold 3 were also generated. To identify the most likely complexes, the top five HDOCK poses showing higher dock score and top five AlphaFold 3 poses showing higher confidence score (Fig. S4, Table S2) were analyzed by MM-GBSA for a comparative binding free energy (Table S3). Based on the higher MM-GBSA score, HDOCK pose-1 and pose-2 were selected for further analysis, however, none of the AlphaFold 3 poses were selected due to their lower MM-GBSA scores. The two HDOCK poses were used for detailed residue-wise interaction analysis between E1-E2 dimer with AGPAT1 to map the critical residues involved in establishing the stable complex. The per-residue decomposition was also carried out to identify the major contributing residues in terms of energetics. We observed that in the pose-2 AGPAT1 interacted with the domain II of the E1 protein (Fig. 3A) that was previously reported to interact with the MXRA8 protein (9). Based on this support from the previous experimental finding, and due to a considerably higher MM-GBSA score (Table S3), the pose-2 was studied further for the complex stability using the molecular dynamics (MD) simulation.

**Figure 3.**
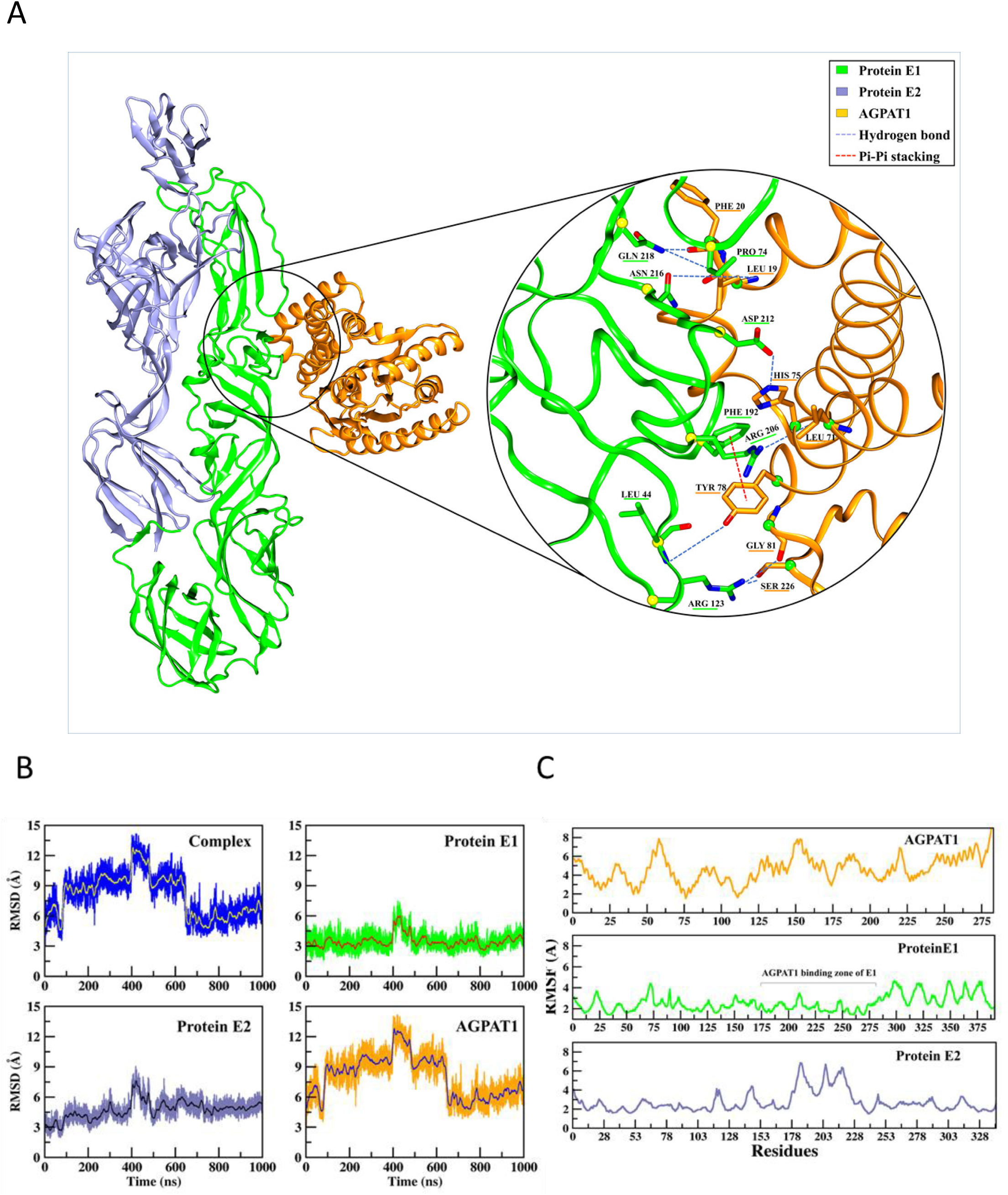
AGPAT1 interaction with the CHIKV E1-E2 dimer *in silico*. (A) The pictorial representation of the CHIKV E1-E2 dimer interaction with the AGPAT1 protein is shown. Interactions in the form of the hydrogen-bonds and Pi-Pi interactions are depicted. (B) The structural conformation of the complex over time was studied as RMSD, calculated using the backbone atoms of the whole complex (E1-E2 dimer with AGPAT1). Individual backbone RMSD of each protein present in the complex is also depicted. (C) The atomic level fluctuation for the individual protein in the complex was studied as RMSF. Significantly less fluctuation in the AGPAT1 binding zone of the E1 protein is highlighted.

### MD simulation of the AGPAT1-CHIKV E1-E2 dimer complex

The MD simulation was carried out to understand the stable state of the docked complex with respect to time. Post-MD analysis techniques, like RMSD and RMSF, were used for the quantitative and qualitative measurements of the MD trajectories. The RMSD study indicated that the complex achieved its stable state in a 1000 ns MD simulation run. The dynamic behaviour of the complex system during the MD simulations revealed that although the structure experienced initial fluctuations, it stabilized after ∼600 ns (Fig. 3B). This suggested that, in the later stages of the simulation, protein-protein interactions contributed to reducing the structural deviations. Further, a comparison between the initial (0 ns) and final (1000 ns) structures of the complex showed a minimal fluctuation (RMSD ∼2.8 Å), indicating the structural robustness and stability of the docked complex.

The individual RMSD profiles of the E1–E2 dimer and AGPAT1 were also evaluated (Fig. 3B). The results indicated that the E1–E2 dimer maintained a stable conformation from the beginning, while AGPAT1 showed some initial fluctuations but stabilizes around ∼600 ns. This suggested that the observed fluctuations in the AGPAT1-CHIKV E1-E2 complex were primarily due to AGPAT1.

The MM-GBSA analysis was carried out on the stable trajectory to embrace the outcomes of the MD simulations. The results indicated that AGPAT1 had a high binding affinity of −85.81 kcal/mol towards the CHIKV E1-E2 dimer. This high binding affinity indicates the robustness of the complex that kept the complex stable during the MD simulation.

Protein flexibility is an intrinsic characteristic that can influence various biological processes. The RMSF analysis (Fig. 3C) showed that the E1-E2 dimer interaction with AGPAT1 reduces the protein flexibility at the binding regions (marked in Fig 3C). Here, AGPAT1 primarily interacted with the E1 component of the E1-E2 dimer to gain high stability with an average Cα RMSF value of ∼2.0 Å, showing a rigid conformation with minimal fluctuation.

### Interaction fingerprinting identifies the key amino acids in the AGPAT1-CHIKV E1-E2 complex

Stable interactions observed between the CHIKV E1–E2 dimer and the AGPAT1 protein throughout the MD simulation were primarily stabilized by strong hydrogen bonds and a notable π–π stacking. Importantly, the majority of the AGPAT1’s contacts were established with domain II of the E1 protein. A total of eight hydrogen bonds (see below) and one π–π stacking interaction (Phe192@Tyr78) were identified as key stabilizing forces during the simulation (Fig 3A). Interaction fingerprint analysis of the most stable conformational state (which is close to the average structure of the lowest RMSD) obtained from the MD trajectory revealed the specific residues involved in the complex formation (Fig. 3A). Within the 4.0 Å distance, AGPAT1 residues closely associated with six E1 residues. These included the hydrophobic (Leu73), aromatic (Phe192, Tyr214), basic (Arg123, Arg206), and polar (Asp212, Asn216, Gln218) amino acids. Among these residues, Arg123, Arg206, Asp212, Asn216, and Gln218 were involved in making stable hydrogen bonds with AGPAT1 residues. The AGPAT1 residues Leu19, Phe20, Ser22, Phe47, Leu71, His75, Tyr78, and Gly81 were in close contact with the E1 protein, contributing to the overall stability of the complex. The key CHIKV E1 and AGPAT1 interactions, which were durable during the MD simulations more than 30% were identified as Phe192@Tyr78, Arg206@Leu71, Leu44@Tyr78, Gln218@Leu19, Asp212@His75, Gln218@Pro74 and Asn216@Leu19.

### CHIKV E1 interacts with AGPAT1 in Huh7 cells

To validate the *in silico* findings, confocal microscopy-based immunofluorescence colocalization studies were done. Colocalization of CHIKV E1 and the endogenous AGPAT1 (PCC 0.63) was observed in CHIKV-infected Huh7 cells (Fig. S5A). Since E1 and E2 may exist as dimers in CHIKV-infected cells, it cannot be reliably inferred that AGPAT1 interacts with CHIKV E1. To address this, Huh7 cells were transfected with the plasmid expressing CHIKV E1 carrying the FLAG tag. Here, we observed the colocalization (PCC 0.69) of the endogenous AGPAT1 with the ectopically-expressed CHIKV E1-FLAG protein (Fig. S5B). To corroborate these findings, Huh7 cells were transfected with the plasmids expressing CHIKV E1-FLAG and AGPAT1-HA proteins. These cells showed strong colocalization (PCC 0.79) of the ectopically-expressed CHIKV E1-FLAG and AGPAT1-HA proteins (Fig. S5C), whereas CHIKV-E2-FLAG did not colocalize (PCC 0.30) with AGPAT1-HA protein (Fig. S5D). These findings were supported by co-immunoprecipitation of CHIKV E1 and AGPAT1 proteins from Huh7 cells (Figs. S4E, S4F).

### The mutants of CHIKV E1 and AGPAT1 proteins show a reduced interaction

The *in silico* interaction analysis followed by the per-residue energy decomposition analysis identified critical residues involved in the AGPAT1 interaction with the E1-E2 dimer (Fig. S6). From these residues, the alanine scanning identified the single and pair-wise amino acids of both proteins, where a change to alanine will disturb the stability of the complex without affecting the protein native structure (Fig. S7). Based on computational interaction energy and interaction type analysis, seven mutations from the E1 subunit and three mutations from AGPAT1 were selected from an initial set of fifteen and six mutations, respectively, for further *in vitro* analysis (Fig. S7). Western blotting established the ectopic expression of the mutant proteins in the transfected Huh7 cells (Figs. 4A, 4D). While CHIKV E1 mutants P74A, R206A, and F192A+R206A showed no difference in their interaction with AGPAT1 as shown by the PLA signals, while mutants F192A, N216A, Q218A, and F192A+N216A showed a significantly reduced (*p*<0.001) interaction compared to the control (Figs. 4B, 4C). For AGPAT1, Y78A and L19A mutations significantly reduced (*p*<0.001) its interaction with CHIKV E1 in the CHIKV-infected cells, while the double mutant L19A+Y78A showed no difference compared to the control (Figs. 4E, 4F). These data confirm the interaction of AGPAT1 with CHIKV E1 and validate the *in silico* findings on critical amino acids involved in the interaction.

**Figure 4.**
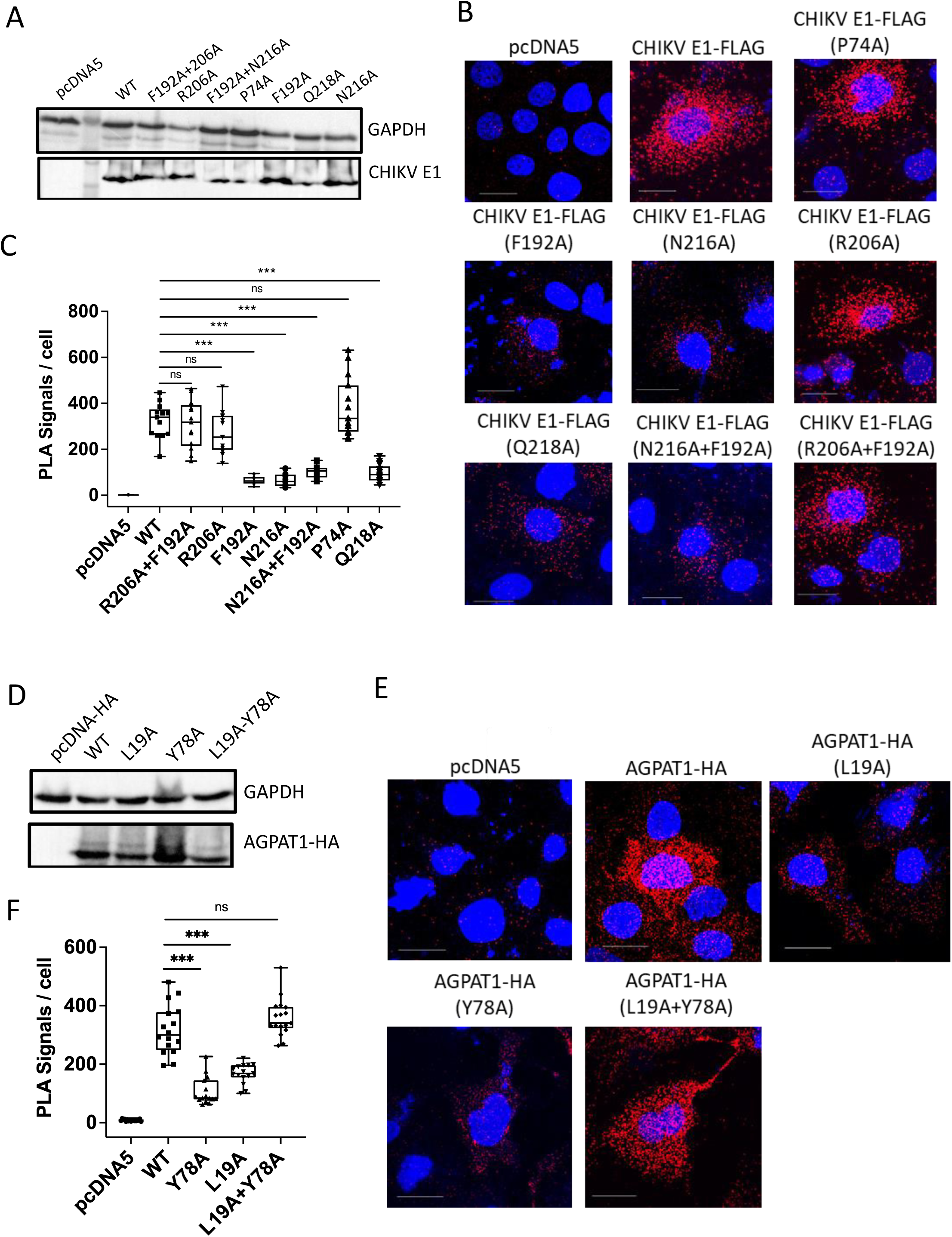
Interaction of the mutants of CHIKV E1 and AGPAT1 proteins. (A) Huh7 cells were co-transfected with pAGPAT1-HA and the plasmids expressing the CHIKV E1-FLAG protein (WT) or its mutants, or the empty vector pcDNA5. The cells were harvested at 48 h pt and Western blotting was done to check the expression of the different CHIKV E1 mutant proteins. (B) The transfected cells were fixed 48 h pt, and the proximity ligation assay (PLA) was performed using the mouse monoclonal anti-FLAG antibody to track the CHIKV-E1-FLAG protein or its mutants, and the rabbit monoclonal anti-HA antibody for the AGPAT1-HA protein. The red dots indicate the interaction of CHIKV E1-FLAG with AGPAT1-HA. The representative confocal microscopy images are shown (scale bar = 25 µm). (C) The PLA signals per cell were determined and shown as the box and whisker plot. The statistical analysis was done by the ordinary one-way ANOVA with Dunnett’s multiple comparisons test considering the WT CHIKV E1 as the control. (D) Huh7 cells were co-transfected with the plasmid expressing the CHIKV E1-FLAG protein and the plasmids expressing the AGPAT1-HA protein (WT) or its mutants, or the empty vector pcDNA5-HA. The cells were harvested 48 h pt and Western blotting was done to check the expression of the different AGPAT1 mutant proteins. (E) The transfected cells were infected 24 h pt, and fixed 24 h post-infection. The PLA was performed using the mouse monoclonal anti-CHIKV E1 antibody to track the CHIKV E1 protein in infected cells, and the rabbit monoclonal anti-HA antibody for the AGPAT1-HA protein (WT) or its mutants. The red dots indicate the interaction of CHIKV E1 with AGPAT1-HA. The representative images of the confocal microscopy are shown (scale bar = 25 µm). (F) The PLA signals per cells were determined and shown as the box and whisker plot. The statistical analysis on data presented at mean±SD was done by the ordinary one-way ANOVA with Dunnett’s multiple comparisons test considering the WT AGPAT1-HA as the control; \*\*\**p*<0.001, ns=not significant.

### CHIKV binding and uptake in Huh7 cells ectopically-expressing the AGPAT1 mutant proteins

CHIKV uptake and replication were studied in Huh7 cells transfected with the plasmids expressing the mutant AGPAT1-HA proteins. The expression of the ectopically-expressed mutant proteins was validated by staining the non-permeabilized cells with HA antibody (Fig. 5A). The CHIKV binding, measured as MIFD on CHIKV-incubated stained cells, was enhanced (*p*<0.001) in the cells expressing the WT AGPAT1-HA (Figs. 5B, 5C). However, this enhancement was lost in the mutants Y78A and L19A. Interestingly, the double mutant Y78A+L19A showed an enhanced (*p*<0.001) CHIKV binding. Along the same lines, CHIKV uptake (Fig. 5D) and viral replication (Fig. 5E) were significantly enhanced in the cells expressing the WT AGPAT1-HA protein. Again, this enhancement was lost in the mutants Y78A and L19A. Notably, the AGPAT1 double mutant L19A+L78A showed an enhanced CHIKV uptake and replication compared to the cells transfected with the pcDNA5 control. These data ratify the earlier finding on protein-protein interaction (Fig. 4) and demonstrate the role of AGPAT1 in CHIKV binding and uptake in Huh7 cells.

**Figure 5.**
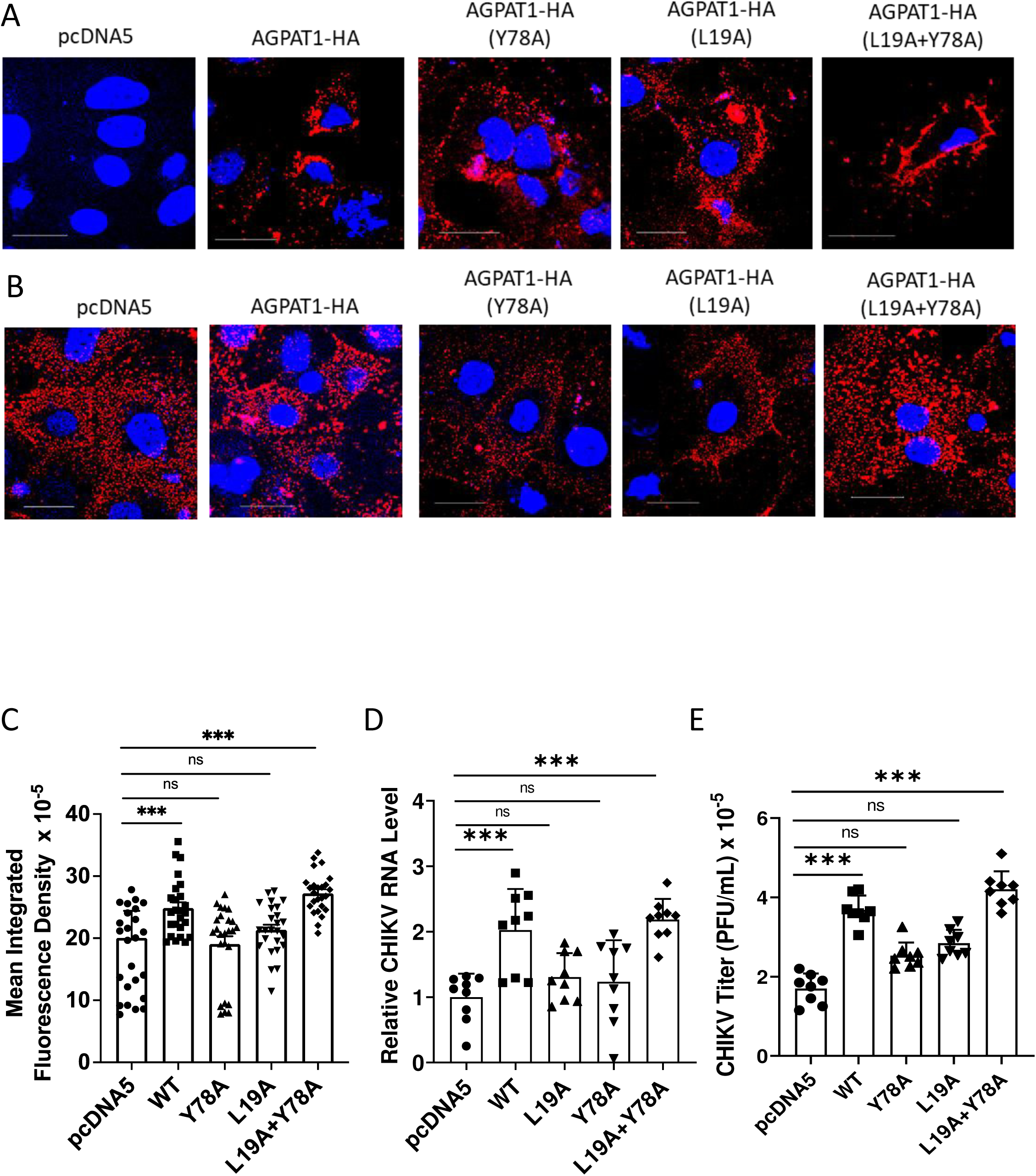
CHIKV binding and uptake in Huh7 cells ectopically expressing the AGPAT1 mutant proteins. (A) Huh7 cells were transfected with the plasmids expressing AGPAT1-HA or the mutant proteins, or the empty vector pcDNA5. The non-permeabilized cells were blocked and then stained with the rabbit monoclonal anti-HA antibody followed by fixing and staining with anti-rabbit Alexa Fluor-568 antibody. The cells were imaged under the Leica SP8 confocal microscope to examine the expression of the ectopically-produced AGPAT1-HA or its mutants on the plasma membrane. The representative images are shown (scale bar = 25 µm). (B) Huh7 cells were transfected with the plasmids expressing AGPAT1-HA or the mutant proteins, or the empty vector pcDNA5. The cells were washed at 48 h pt and incubated with CHIKV (MOI 50) on ice for 1 h to allow the virus binding. Cells were then washed with ice-cold PBS, blocked with 5% BSA, and stained with the monoclonal mouse anti-CHIKV E2 antibody followed by anti-mouse Alexa-647 secondary antibody to visualize the presence of the CHIKV particles on the plasma membrane surface. The cells were imaged under the Leica SP8 confocal microscope (scale bar = 25 µm). The representative images are shown. (C) The MIFD data of the transfected Huh7 cells stained as above for the CHIKV binding are presented from 3 biological replicates. (D) Huh7 cells were transfected with the plasmids expressing AGPAT1-HA or the mutant proteins, or the empty vector pcDNA5. The cells were incubated 48 h later with CHIKV (MOI 3) for 1 h on ice followed by a 37 °C incubation for 1 h for uptake. The cells were treated with trypsin to remove the extracellular virion particles before harvesting. The cells were lysed to isolate the RNA. The CHIKV RNA in the cells was quantified by qRT-PCR. The relative CHIKV RNA levels are presented where the CHIKV RNA level in cells transfected with pcDNA5 was taken as 1. The data from 5 biological replicates and 2 technical replicates are shown. (E) Huh7 cells were transfected with the plasmids expressing AGPAT1-HA or the mutant proteins, or the empty vector pcDNA5. The cells were infected 48 h later with CHIKV (MOI 3) and incubated at 37 °C. The virus titers in the culture supernatant at 6 h pi are presented. The data from 4 biological replicates and 2 technical replicates are shown. The statistical analysis on data shown as mean±SD was done by the one-way ANOVA with Dunnett’s multiple comparisons test considering the pcDNA5-transfected cells as the control; \*\*\**p*<0.001, ns=not significant.

### Role of AGPAT1 in binding and uptake of Ross River virus (RRV)

RRV, an alphavirus, shows significant conservation of the E1 amino acids involved in CHIKV E1 binding with AGPAT1 (Fig. 6A). However, Sindbis virus (SINV), another alphavirus, shows little conservation of these E1 residues. Interestingly, RRV binding to AGPAT1 KO HAP1 cells was reduced by 85% compared to the WT cells (*p*<0.01), whereas SINV binding was not affected (Fig. 6B). Similarly, RRV uptake in the KO cells was lower (*p*<0.01) by 56% than in the WT cells; however, SINV uptake was not affected (Fig. 6C).

**Figure 6.**
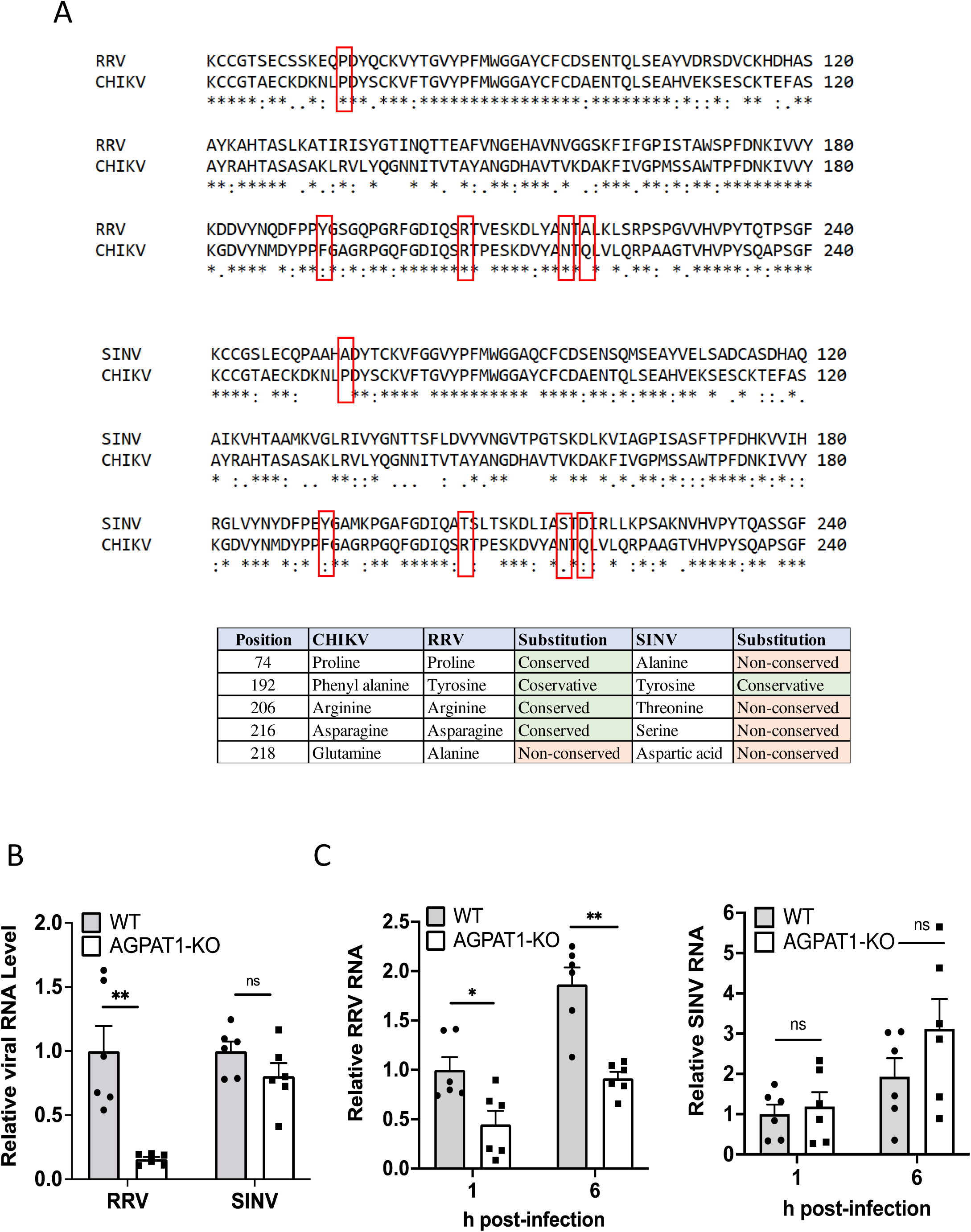
Role of AGPAT1 in RRV binding and uptake in Huh7 cells. (A) The amino acid sequence of the CHIKV E1 protein (GenBank ID JF274082) was aligned with the RRV E1 protein (GenBank ID GQ433359.1) and SINV E1 protein (GenBank ID V01403.1) using the program Clustal Omega. Amino acid numbering is provided at the right. Conserved amino acids are marked with an asterisk (*), conservative changes as (:), whereas semi-conservative changes are marked as (.) under the aligned amino acid sequence. The amino acids involved in CHIKV E1 interaction with AGPAT1 are boxed. The table lists the amino acid substitutions between the CHIKV E1, RRV E1, and SINV E1 proteins and identifies the nature of the change. (B) The KO and WT HAP1 cells were incubated with RRV and SINV on ice for 30 min at 5 MOI to allow virus binding. The unbound virus was removed by ice-cold PBS wash. The cells were then lysed, and total RNA was extracted. The virus binding to cells was determined by the level of the viral RNA using qRT-PCR. The relative viral RNA levels are presented where the viral RNA level in the WT cells was taken as 1. The data from 3 biological replicates and 2 technical replicates are shown. (C) The WT and AGPAT1 KO HAP1 cells were incubated with RRV or SINV (MOI 1) for 1 h on ice. The cells were washed with chilled PBS before incubating the cells at 37 °C; the cells were treated with trypsin before harvesting to remove the extracellular virion particles. The cells were harvested at the indicated times to isolate the total RNA. The viral RNA in the cells was quantified by qRT-PCR. The relative viral RNA levels are presented where the RNA level in the WT cells at 1 h was taken as 1. The data from 3 biological replicates and 2 technical replicates are shown. The statistical analysis on data presented as mean±SD used the Student’s t-test with Welch’s correction; \**p*<0.05, \*\**p*<0.01, ns=non-significant.

### Role of AGPAT1 in CHIKV binding in ERMS cells

CHIKV replication in skeletal muscle cells is a critical mediator of CHIKV pathogenesis (34–38). We, therefore, studied the role of AGPAT1 in CHIKV infection in ERMS cells of the human skeletal muscle lineage. Here also, AGPAT1 was present on the plasma membrane (Fig. S8). AGPAT1 antibodies caused ∼70% reduction (*p*<0.001) of CHIKV binding to ERMS cells based on MIFD, suggesting that CHIKV uses AGPAT1 to bind to ERMS cells also.

## DISCUSSION

Virus receptors determine the host specificity and cell tropism, and the knowledge of the virus-host receptor interactions could help develop novel antivirals targeting the very first step in the virus infection of the host cell. So far, Mxra8, Prohibitin, and CD147 proteins have been described as CHIKV receptors in mammalian hosts. Of these, the Mxra8 receptor for CHIKV has been extensively characterized (9–11, 27, 39–41). Mxra8 had a role in CHIKV infection in mice and several human cell lines (23, 27). However, cells such as HEK-293T and Huh7 (27) or HAP1 (24), lacking the surface expression of MXRA8, are susceptible to CHIKV infection, suggesting that additional host proteins serve as CHIKV receptors in mammalian cells. We have identified AGPAT1 as a CHIKV receptor on human cell lines such as Huh7, ERMS, and HAP1. Importantly, these cells lack MXRA8 expression on their plasma membrane (Fig. S9). Thus, the identification of AGPAT1 as an additional receptor is a significant advancement in understanding the CHIKV pathogenesis in humans, especially in the context of CHIKV infection of cells lacking MXRA8.

AGPAT1 is an enzyme involved in lipid metabolism (42), the development of skeletal muscle (43), and the physiology of multiple organ systems (44). AGPAT1 has not been reported to have a direct role in the virus life cycle, although its involvement in lipid metabolism can indirectly influence viral replication (45, 46). Thus, this is the first report showing a direct role for AGPAT1 in the virus life cycle as a novel CHIKV receptor on mammalian cells.

Different methods have been employed to study the CHIKV receptor. For example, Prohibitin was identified as binding to CHIKV using the ‘Virus Overlay Protein Binding Assay’ (VOPBA) on human microglial CHME-5 cells (28). On the other hand, Mxra8 was identified by CRISPR-Cas9 screening for CHIKV infection in the mouse fibroblast NIH3T3 cells (27). The Strep-tagged CHIKV envelope protein complex was used to identify CD147 protein in HEK-293T cells (29). We employed a pull-down method to detect the Huh7 plasma membrane proteins directly binding to CHIKV virions. The mass spectrometry identified several proteins, including Prohibitin, which validated the method used in our study. However, we did not detect MXRA8 in our experiments. This may be related to the low level of MXRA8 or its absence in the Huh7 cell membrane (Fig. S9) compared to the mouse fibroblast NIH 3T3 cells where Mxra8 was identified as a CHIKV receptor (27). The CRISPR-Cas9 screening for CHIKV infection in the mouse fibroblast NIH3T3 cells did not pick up Prohibitin or Agpat1 (27). For AGPAT1, this may be related to its absence on the NIH3T3 cell membrane (Fig. S9).

AGPAT1 is primarily located in the ER membranes (33). For AGPAT1 to act as a virus receptor, it must be located on the plasma membrane. Our study demonstrated the presence of AGPAT1 on the plasma membrane of Huh7, HAP1 and ERMS cells by confocal microscopy on non-permeabilized cells and Western blotting of the plasma membrane fraction of Huh7 cells. In the mouse myoblast C2C12 cells (Fig. S9) also, AGPAT1 was found on the plasma membrane as reported before (43).

Using the pull-down-mass-spectrometry method, we identified AGPAT1 from Huh7 cells as a protein binding with CHIKV virions. The interaction of AGPAT1 with CHIKV was verified by co-immunoprecipitation, while the binding of CHIKV with AGPAT1 was demonstrated on Huh7, HAP1, and ERMS plasma membrane by confocal microscopy and other methods. Antibody to AGPAT1 inhibited CHIKV binding to the plasma membrane and reduced virus uptake in Huh7 and ERMS cells. The role of AGPAT1 in CHIKV binding and uptake was further confirmed in HAP1 cells, where CHIKV binding, uptake, and replication were significantly reduced in AGPAT1 KO cells. Ectopic expression of AGPAT1 in the KO cells rescued the reduced CHIKV binding, uptake, and replication; a huge enhancement (2300%) of CHIKV uptake was seen in the rescued cells. Further, the ectopic expression of AGPAT1 on Huh7 plasma membrane enhanced CHIKV binding to cells and this enhancement was not seen with AGPAT1 mutants. These data demonstrate that AGPAT1 has an important role in CHIKV binding and uptake in Huh7, HAP1, and ERMS cells.

CHIKV is an enveloped virus projecting the viral envelop proteins E1 and E2 on its outer surface (47). The heterodimer of the E1 and E2 proteins is present as a trimer on the virion envelope (47, 48). Our *in silico* studies predicted the interaction of AGPAT1 with the CHIKV E1-E2 dimer at the E1 surface. This was validated by confocal microscopy, showing the colocalization of CHIKV E1 protein with AGPAT1 in Huh7 cells. The *in silico*-predicted critical amino acids at the E1 surface majorly interacting with AGPAT1 were validated using the mutant proteins. While no information is available on how Prohibitin interacts with CHIKV, mammalian MXRA8 was shown to interact with CHIKV E1 as well as E2 protein (9). Here, we have demonstrated AGPAT1 interaction with CHIKV E1 and identified the key amino acids of the proteins involved. Interestingly, some of these residues of CHIKV E1 were also engaged in CHIKV interaction with mammalian MXRA8 (9). It may be noted that the avian MXRA8 was shown to interact with the E1 protein of Western equine encephalitis virus, yet another alphavirus (11).

CHIKV has an extensive host range, infecting cells of multiple origins in human and mosquito hosts. It is, therefore, likely that the virus uses as receptor a protein ubiquitously present on cells of different origins. Alternatively, the virus may use different proteins on different cells. The latter seems to be the case as AGPAT1, Mxra8, Prohibitin, and CD147 have been identified as CHIKV receptor proteins on different cell types. Again, these may not be the only proteins the virus uses as a receptor, and, therefore, the search for additional proteins as CHIKV receptors must continue.

## MATERIALS AND METHODS

### Cells and viruses

The Vero, BHK-21, Huh7, NIH 3T3, and HeLa cell lines were obtained from the National Centre for Cell Sciences (NCCS) cell repository. The human embryonal rhabdomyosarcoma (ERMS) (RD-CCL-136-ATCC), and C2C12 (CRL-1772) were obtained from the ATCC, USA. The minimum essential medium, Eagle (MEM) (HiMedia, India; AL0475), was used to cultivate Vero and BHK-21 cells. Dulbecco’s modified Eagle medium (DMEM) (HiMedia, India; AL007A) was used to culture Huh7, ERMS, and C2C12 cells. HAP1 cells were cultured using Iscove’s modified Dulbecco’s Medium (IMDM) (HiMedia, India; AL070S). All the media were supplemented with 10% fetal bovine serum (FBS) (Gibco; 10270106) and 1X penicillin-streptomycin-glutamine (PS) solution (HiMedia, India; A001A). The cells were grown at 37 °C under a 5% CO_2_ atmosphere.

The IND-06-Guj (JF274082; GenBank) was used in this study. Ross River and Sindbis viruses were obtained from the World Reference Center for Emerging Viruses and Arboviruses, The University of Texas Medical Branch, USA.

### Purification of Chikungunya virus

The sucrose gradient purified CHIKV was used in this study. BHK-21 cells were cultured in a T175 flask, infected with CHIKV for 1 h at 0.1 MOI, and incubated at 37 °C for 36 h. The culture supernatant was collected, and the virus was precipitated by 8% PEG-8000 at 4 °C. The virus pellet was suspended in the NTE buffer (150 mM NaCl, 50 mM Tris, and 5 mM EDTA, pH 8.0) and over-layered on a discontinuous 30-60% sucrose gradient. The virus was ultracentrifuged overnight at 4 °C in a SW28Ti rotor (Beckman Coulter) at 26,000 rpm. The virus layer collected from the gradient interphase was over-layered on a 20% sucrose bed (2 ml) and pelleted at 4°C for 6 h using SW41Ti rotor (Beckman Coulter) at 39,000 rpm. The virus pellet was re-suspended in MEM and stored in small aliquots at −80 °C. The virus purity was checked by SDS-PAGE and its titer was determined by plaque assay on Vero cells.

### Plaque assay for CHIKV titration

Vero cells were plated overnight in a 6-well plate to obtain 70% confluency. Serial dilutions of the CHIKV sample were made in MEM, and 200 µl virus from different dilutions was used to infect the cells at 37 °C for 1 h. The wells were then washed and overlaid with 1% agarose in MEM. The plates were incubated at 37 °C for 36 h to obtain the plaques. The cell monolayers were fixed with 4% formalin for 2 h and stained with 0.2% crystal violet for 1 h. The plates were then washed with water and air-dried. The plaques formed were counted, and the virus titer was calculated as plaque-forming unit (PFU)/ml using the dilution factor.

### Purification of the plasma membrane proteins

The Huh7 cell monolayer grown in T175 flasks was washed with 1X PBS and scrapped in the same. Dounce Homogenizer (100 cycles) was used to lyse the cells with the homogenization buffer containing the protease inhibitor cocktail. The plasma membrane proteins were isolated using the Plasma Membrane Isolation Kit (ab65400; Abcam). The total cell lysate and the cytosolic and plasma membrane protein fractions were run on an SDS-PAGE gel, and Western blotted. E-cadherin was used as a marker for the plasma membrane proteins fraction, GAPDH as a marker for the cytoplasmic proteins, and Calreticulin was used to detect the Endoplasmic Reticulum (ER) membrane contamination in the preparation. The following antibodies were used for the Western blotting: rabbit monoclonal anti-E-Cadherin antibody (CST 3195; Cell Signalling Technologies), rabbit polyclonal GAPDH antibody (GTX100118; GeneTex), rabbit polyclonal anti-Calreticulin antibody (2891; Cell Signalling Technologies), and mouse polyclonal AGPAT1 antibody (ab67018; Abcam).

### Pull-down of plasma membrane proteins with the agarose-conjugated CHIKV

The sucrose gradient purified CHIKV (150 µg) was immobilized on Amino-Link™ agarose beads (Thermo Fisher Scientific) by amine linkage chemistry using sodium cyanoborohydride. The Pierce^TM^ co-immunoprecipitation kit (Thermo Fisher Scientific) was used to incubate the CHIKV beads with the Huh7 plasma membrane proteins in 1X PBS, 5% octyl-β-D-glucopyranoside. The beads were washed with 1X PBS, 5% octyl-β-D-glucopyranoside, and the bound proteins were eluted using a low pH buffer (Glycine, pH 2.8) followed by neutralization with 1.5 M Tris pH 9. The eluted proteins were run on an SDS PAGE, followed by the gel extraction. The proteins were trypsin (V5280, Promega) digested and subjected to mass spectrometry using an Electrospray Ionization-Mass Spectrometer: SCIEX TripleTOF^®^ 5600+. The mass spectrometry output was analyzed through MASCOT and Protein Pilot to identify the proteins.

### Immunostaining for localization of AGPAT1 on plasma membrane

The cells were grown overnight at 37 °C on coverslips (Bluestar) placed in a 24-well plate (Corning). The coverslips were washed with chilled 1X PBS and blocked with 5% BSA on ice. For localization of AGPAT1, the cells were incubated on ice for 1 h with rabbit polyclonal anti-AGPAT1 (ab235328; Abcam). The cells were then fixed with 2% formaldehyde for 20 min at room temperature, followed by 30 min incubation of anti-rabbit Alexa Fluor-568 antibody (A11011; Invitrogen) for staining AGPAT1. The coverslips were mounted on Prolong Gold antifade with DAPI (P36935; Invitrogen) and observed under the Leica SP8 confocal microscope.

### Immunostaining of proteins in the permeabilized mammalian cells

The cells grown overnight at 37 °C on coverslips (Bluestar) placed in a 24-well plate (Corning) were washed twice with PBS and fixed with 2% paraformaldehyde for 20 min at room temperature. For permeabilization, the cells were incubated with 0.3% Tween-20 or 4% paraformaldehyde and blocked with 5% BSA for 1 h at room temperature. The cells were stained for 1 h at room temperature with rabbit polyclonal anti-AGPAT1 antibody (ab235328; Abcam) for AGPAT1, mouse monoclonal anti-E1 (GTX135187; R&D systems) for CHIKV E1, rabbit monoclonal anti-HA (3724; Cell Signalling Tech) or mouse monoclonal anti-HA (2367; Cell Signalling Tech) for AGPAT1-HA, rabbit monoclonal anti-FLAG (14793; Cell Signalling Tech) or mouse monoclonal anti-FLAG (91878; Invitrogen) for E1-FLAG. After washing with PBS, the cells were incubated at room temperature for 1 h with anti-mouse Alexa Fluor-647 antibody (4410S; Cell Signalling Tech), anti-rabbit Alexa Fluor-568, or anti-rabbit Alexa Fluor-488 antibody (A11008; Invitrogen). The coverslips were mounted with Prolong Gold antifade with DNA stain DAPI (Thermo Fisher Scientific) and observed under the Leica SP8 confocal microscope. The images were processed using LasX software (Leica).

### siRNA-mediated knockdown of AGPAT1

The Mission® esiRNA (EHU228041; Merck), containing a pool of siRNAs targeting the human AGPAT1 mRNA, was used to knockdown the AGPAT1 protein expression in Huh7 cells. The Mission® esiRNA Universal Negative control #1 (SIC001; Merck) containing the non-targeting pool of siRNAs was used as the control. The esiRNAs at 30 nM concentration were transfected into Huh7 cells using Lipofectamine RNAiMAX reagent (13778075; invitrogen). At 48 h post-transfection (pt), the knockdown of the protein was quantified by Western blotting. GAPDH was used as the control.

### Ectopic expression of protein

The cDNA encoding AGPAT1 or its mutants fused with the HA-tag sequence were cloned in pcDNA5 vector under the CMV promoter. The primers used for the cDNA amplification are described in Table S1. The cDNA encoding CHIKV E1 or its mutants fused with the FLAG-tag sequence were cloned in pcDNA5 vector under the CMV promoter. The cells were transfected with the expression plasmids using Lipofactamine 3000 transfection reagent (L3000015; Invitrogen). The ectopic expression of the protein was verified by Western blotting of the cell lysate with an appropriate antibody. GAPDH was used as the loading control.

### Total RNA isolation, reverse transcription, and quantitative PCR

The total RNA was isolated from the cells using RNAiso Plus (TaKaRa), precipitated by absolute isopropanol, air-dried, and reconstituted in nuclease-free water. The purity and concentration of RNA were determined spectrophotometrically. The cDNA was made using the ImProm-II reverse transcription kit (Promega), which used 50-500 ng RNA and random primers. The cDNA was quantified by qPCR using SYBR^®^ green chemistry. The TB Green Premix Ex Taq II (Tli RNaseH Plus) (TaKara) was used as the PCR master mix. The following CHIKV nsP2 primers were used to detect the CHIKV RNA: Forward- GGCAGTGGTCCCAGATAATTCAAG, Reverse- GCTGTCTAGATCCACCCCATACATG. The RRV RNA was detected by primers for E1: Forward- GCGACGGT GGATGTCAAGGAG, Reverse- AGCCAGCCCACCTAACCCACTG. The SINV RNA was detected using the following primers for E2: Forward- AAAGGATACTTTCTCCTCGC, Reverse- TGGGCAACAGGGACCATGCA. The human GAPDH RNA was detected by the GAPDH forward primer GGTGAAGGTCGGAGTCAACG and GAPDH reverse primer AGGGATCTCGCTCCTGGAAG. The qPCR was run in the standard curve mode with a melting curve plot under the SYBR green chemistry in a Quantstudio^TM^ 6 Flex (Applied Biosystems) PCR machine. The data obtained were normalized using the cellular GAPDH levels.

### CHIKV binding assay

For CHIKV localization and binding studies, the cells were incubated for 1 h with CHIKV on ice, followed by blocking with 5% BSA and 1 h incubation with mouse monoclonal anti-E2 (MAB1219; Native Antigen) in 1% BSA on ice. The cells were then fixed with 2% formaldehyde for 20 min at room temperature, followed by 30 min incubation of anti-rabbit Alexa-568 antibody (A11011; Invitrogen) for staining AGPAT1, or anti-mouse Alexa Fluor-647 antibody (4410S; Cell Signalling Technologies) for staining the CHIKV E2 protein. The coverslips were mounted on Prolong Gold antifade with DAPI (P36935; Invitrogen) and observed under the Leica SP8 confocal microscope. The images were processed by the LasX software (Leica). To determine the mean fluorescence intensity density (MFID), five different fields with an ROI (Region of Interest) of fixed area, having at least 10 cells, were taken into account. The mean of pixel densities from 5 middle planes were taken from each field before calculating the mean of all the fields.

### CHIKV uptake and replication

The virus uptake was studied by determining the CHIKV RNA levels early during the infection. To this end, the cell monolayer was incubated with CHIKV for 1 h on ice followed by wash with chilled PBS. Complete media was added to the cells and incubated at 37°C. The culture supernatants were collected and the cells were treated briefly with trypsin solution (TCL007; HiMedia, India) before harvesting at different times points post-infection. The total RNA was isolated from the cells and the virus titers were determined in the culture supernatant. The CHIKV RNA levels were determined by qRT-PCR and the viral titers were determined by plaque assay. The viral RNA level recorded at 1 h pi was taken as the indicator of the virus uptake and used to calculate the enhancement/reduction in the virus uptake.

### Generation of the AGPAT1 knockout HAP1 cell line

The AGPAT1 knockout (KO) HAP1 cells were generated using the CRISPR-Cas9 technology. The sgRNA sequence 5’-CTCAGCATCAAAGTTAGTAT-3’ was designed using the Broad Institute database for sgRNAs and the CHOPCHOP online predictor, and obtained from Genscript as incorporated inside the shuttle vector pSpCas9 BB-2A-Puro (PX459) V2.0. The plasmid containing the sgRNA was transfected into HAP1 cells that were subjected to puromycin selection at 1 µg/ml for 6 passages. The stable cells were subjected to the single-cell clonal selection to obtain a pure AGPAT1 KO cell line. Western blotting was done to establish the absence of the AGPAT1 protein in the KO cells.

### Ectopic expression of AGPAT1 and CHIKV E1 proteins

The ectopic expression of the AGPAT1 and CHIKV E1 proteins was obtained by transfecting the cells with the respective expression plasmid. The expression plasmids were made by inserting the respective cDNAs into the expression vector pcDNA5 (Thermo Fisher Scientific) under the CMV promoter. The plasmid nucleotide sequence was confirmed by Sanger sequencing. The AGPAT1 protein was expressed as a fusion protein with the HA tag, and the CHIKV E1 protein was fused with the FLAG tag. The following primers were used for amplification of the AGPAT1-HA cDNA: Forward, TCA<u>AAGCTT</u>GCCACCATGGATTTG TGGCCAGGGGC (HindIII site underlined); Reverse, CAGT<u>CTCGAG</u>TTA*TGCATAATC CGGAACATCATACGGATA*CCCACCGCCCCCAGGCTTCT (XhoI site underlined, HA-tag sequence in italic). The following primers were used for amplification of the CHIKV E1-FLAG cDNA: Forward, ACC<u>AAGCTT</u>GCCACCATGGGCTACGAACACGTAACAGT GATCCCGAACAC (HindIII site underlined); Reverse, TCA<u>GGATCC</u>TTA*CTTATCGT CGTCATCCTTGTAATC*GTGCCTGCTGAACGACACGCATAGC (BamHI site underlined, FLAG-tag sequence in italic).

### System preparation for the *in silico* studies

The 3D structure of AGPAT1 was taken from the AlphaFold database and CHIKV E1-E2 dimer structure was retrieved from the protein data bank (PDB 3N42)(9, 49). To prepare the protein structures, Maestro’s Protein Preparation Wizard module with OPLS4 force field was used; the protein’s missing regions were filled by the prime module, and the preliminary minimization was done by OPLS4 force field. The systems were prepared with the protonation states of the residues at neutral pH as predicted by the PROPKA module in the Protein Preparation Wizard (50, 51).

### Protein-protein docking

The protein-protein docking of the CHIKV E1-E2 dimer with AGPAT1 was performed using the HDOCK server and AlphaFold 3. A total of 100 docked poses obtained from HDOCK, and top 5 poses of AlphaFold 3 were taken for analysis. The best poses from the docked models were chosen on the basis of docking score, docking confidence and cluster size.

### Molecular dynamics (MD) simulation

The most likely pose obtained from the protein-protein docking was used for the MD simulation using the “Desmond v6.1” module of the Schrodinger suite. The complex was firstly minimized with the OPLS4 force field and then solvated with a predetermined TIP3P water solvent model (52). The size of the repeating unit buffered at 12.0 Å distances was determined by placing them all in the orthorhombic periodic boundary conditions (53, 54). Using a steepest-descent integrator for 2000 steps, the complex underwent energy minimization for 500 ps. For the NPT ensemble simulations, the Nose-Hover chain thermostat and the Martyna-Tobias-Klein barostat were assigned. The RESPA integrator was utilized with a time step of 0.002 ps. The cutoff radius for the short-range Coulombic interactions was 9.0. Bonds were restricted to hydrogen by using the M SHAKE algorithm of DESMOND and the final production run was conducted for one microsecond simulations; the coordinates were stored every 10 ps. The data from the MD simulation trajectory was post-processed using the Schrodinger and VMD scripts (55). Various measurements were obtained to analyze the protein’s structural behaviour from the quantitative analysis including RMSD, RMSF, and MM-GBSA. These measurements provided information about the backbone, side chains and protein-ligand contacts (50, 51, 56–58).

### Binding Free energy calculations

The Molecular Mechanics Generalized Born Surface Area (MM-GBSA) method was employed for the binding free energy calculations using the PRIME module (59). To determine the average binding energy, the snapshots were extracted from the last stable trajectory for analysis. The binding energy (ΔGbind) was computed using the following equation: ΔGbind = ΔE_MM + ΔG_solv + ΔG_SA. Here, ΔE_MM represents the difference in minimized energies between the protein-protein complex, ΔG_solv corresponds to the difference in GBSA solvation energy between the complexes and the sum of the solvation energies of the CHIKV E1-E2 dimer and AGPAT1, and ΔG_SA denotes the difference in surface area energy between the complex and the sum of the surface area energies of the E1-E2 dimer and AGPAT1 (55). The MM-GBSA approach, together with per-residue energy breakdown analysis, allowed us to determine the energy contributions of individual amino acids to identify critical residues at the interface and to reveal primary residue interactions within the complex by decomposing binding free energy (60).

### Prediction of the critical mutations

The goal of the mutational study was to understand the effect of certain point mutations on the interaction of the CHIKV E1-E2 dimer with AGPAT1. To this end, residue scanning in BioLuminate (61) was carried out using the biologics residue scanning panel, using sidechain prediction, and backbone minimization as refinement options and a cutoff distance of 0.0. BioLuminate calculated the net change in protein stability due to the mutation using the prime energy function, which included an implicit solvent term. Stability is defined as the free energy difference between the mutant and natural states caused by a single point mutation. The Schrodinger module BioLuminate (61) was used to calculate the energy using the MM-GBSA method, with force field OPLS4 and solvent model VSGB (58).

### Site-directed mutagenesis

The site-directed mutagenesis was done by polymerase chain reaction (PCR) using the overlapping primers on the plasmids AGPAT1-HA and CHIKV E1-FLAG. Overlapping primers were designed according to the required changes in the coding region. The PCR was done by Phusion® polymerase (NEB, M0530S) using 5X HF buffer; the primers designed for the sites are mentioned in Table S1. The PCR product was treated with DpnI (NEB, R0176S) to digest the methylated plasmid (the template), and the resultant product was PCR purified by a PCR purification kit (Qiagen, 28506). The purified DNA was used to transform *E. coli* DH5α cells. The bacteria were grown for 16 h at 37 °C, and the plasmid DNA was obtained. Sanger sequencing of the plasmid DNA was used to verify the desired mutation.

### Proximity Ligation Assay

The Duolink *in situ* Red PLA kit (Sigma Aldrich; DUO92101-1KT) was used following its recommended protocol. Briefly, Huh7 cells were co-transfected with the plasmids or transfected with a plasmid and infected with CHIKV. The cells were washed, fixed, permeabilised, blocked, and incubated with the primary antibodies for 2 h at room temperature. The primary antibodies used were mouse anti-AGPAT1 antibody and rabbit anti-FLAG or rabbit anti-CHIKV E1 antibodies. The rabbit plus and mouse minus probes were used for the rolling circle amplification. The coverslips were mounted in Duolink *in situ* mounting medium with DAPI and imaged under a confocal microscope. ImageJ software quantified the acquired images by measuring the PLA signals per cell.

## FIGURE LEGENDS

**Figure S1. Identification of CHIKV-binding plasma membrane proteins.**

Purified CHIKV conjugated with agarose beads was incubated with the plasma membrane proteins from Huh7 cells. The pulled-down proteins were identified by mass spectrometry. The figure shows the proteins identified two or more times from the six independent experiments. The NCBI IDs of the proteins are shown on the left.

**Figure S2. AGPAT1 is present on the plasma membrane of Huh7 cells.**

(A) Huh7 cells were permeabilized with 0.3% Tween-20 or were not permeabilized. The cells were then incubated with rabbit AGPAT1 antibody followed by fixing the cells in 2% paraformaldehyde and staining with Alexa Fluor-568 antibody. The nuclei were stained with DAPI, and the cells were imaged using the Leica SP8 confocal microscope (scale bar = 25 µm). The representative images are shown. (B) Huh7 cells were grown as monolayers and the total cell lysate (L) was prepared. The plasma membrane (M) and cytoplasmic (C) fractions of the cells were prepared and Western blotted. The presence of AGPAT1 in the membrane fraction is seen. E-cadherin, GAPDH, and calreticulin were used as controls to demonstrate the protein fractionation.

**Figure S3. CHIKV uptake in Huh7 cells is related to the AGPAT1 protein levels.**

(A) Huh7 cells were transfected with siRNA to AGPAT1 (siAGPAT1) or a control non-targeting siRNA (siNT) and incubated for 48 h at 37 °C. Western blotting showed a ∼60% reduction in the AGPAT1 levels in the siAGPAT1-treated cells. A representative blot is shown. (B) The siRNA-treated cells were incubated 48 h post-transfection (pt) with CHIKV (MOI 1) for 1 h on ice. The cells were then washed with ice-cold PBS and incubated at 37 °C for viral uptake. The cells were treated with trypsin to remove the extracellular virion particles before harvesting. The cells and culture supernatants were harvested at the indicated times. The total RNA was isolated from the cells and the virus titers were determined in the culture supernatant. The CHIKV RNA levels were determined by qRT-PCR (left panel). The level of CHIKV RNA in the siNT-transfected cells at 1 h pi was taken as 1. The viral titers determined by plaque assay are shown in the right panel. The data from 4 biological replicates and 2 technical replicates are shown. (C) Huh7 cells were transfected with the pAGPAT1 plasmid expressing the HA-tagged AGPAT1 or the empty vector pcDNA5. The cells were harvested at 24 and 48 h later. The cell lysates were Western blotted with monoclonal anti-HA antibody to detect the expression of the HA-tagged AGPAT1. GAPDH was used as the loading control. A representative blot is shown. (D) Huh7 cells were transfected with pAGPAT1-HA or pcDNA5, and 48 h later incubated with CHIKV (MOI 5) for 1 h on ice. The cells were washed with ice-cold PBS and then incubated at 37 °C. The cells and the culture supernatants were harvested at the indicated times. The CHIKV RNA in the cells was quantified by qRT-PCR and the viral titers in the culture supernatant were determined by plaque assay. The relative CHIKV RNA levels are presented (left panel), where the CHIKV RNA level at 1 h pi was taken as 1. The right panel has the virus titers. The data from 3 biological replicates and 2 technical replicates are shown. The statistical analysis on data presented as mean±SD was done using the Student’s t-test with Welch’s correction; \**p*<0.05 \*\**p*<0.01, \*\*\**p*<0.001.

**Figure S4: The top poses obtained from the protein-protein docking.**

The tope 5 poses of AGAPAT1 interaction with the CHIKV E1-E2 dimer predicted using the HDOCK and AlphaFold 3 methods are presented. The top poses of AGPAT1 in the complexes (from pose-1 to pose-5) are depicted in grey, orange, pink, blue and red, respectively. The CHIKV E1 in shown in green and E2 in ice blue colour.

**Figure S5. CHIKV E1 interacts with AGPAT1 in Huh7 cells.**

(A) Huh7 cells were infected with CHIKV (MOI 5), and 24 h later were fixed and permeabilized. The cells were stained with rabbit anti-AGPAT1 polyclonal antibody and mouse CHIKV anti-E1 monoclonal antibody, followed by incubation with anti-mouse Alexa Fluor-647 and anti-rabbit Alexa Fluor-488 antibodies. The nuclei were stained with DAPI, and the cells were imaged using the Leica SP8 confocal microscope (scale bar = 20 µm). The PCC was determined for the colocalization of Alexa Fluor-488 (AGPAT1) and Alexa Fluor-647 (CHIKV E1) dyes. The right panel shows the line plot showing the colocalization of the red (CHIKV E1) and green (AGPAT1) fluorescence signals. (B) Huh7 cells were transfected with the plasmid expressing the tagged protein CHIKV E1-FLAG (pE1-FLAG), and 48 h later were fixed and permeabilized. The cells were stained with rabbit anti-AGPAT1 polyclonal antibody and mouse anti-FLAG monoclonal antibody, followed by incubation with anti-mouse Alexa Fluor-488 and anti-rabbit Alexa Fluot-568 antibodies. The nuclei were stained with DAPI, and the cells were imaged using the Leica SP8 confocal microscope (scale bar = 20 µm). The PCC was determined for the colocalization of the Alexa Fluor-568 (AGPAT1) and Alexa Fluor-488 (CHIKV E1) dyes. The right panel shows the line plot showing the colocalization of the green (CHIKV E1) and red (AGPAT1) fluorescence signals. (C) Huh7 cells were transfected with the plasmids pE1-FLAG and pAGPAT1-HA. The cells were fixed 48 h later, permeabilized, and stained with rabbit anti-HA and mouse anti-FLAG antibodies. This was followed by incubation with anti-mouse Alexa Fluor-488 and anti-rabbit Alexa-568 antibodies. The nuclei were stained with DAPI, and the cells were imaged using the Leica SP8 confocal microscope (scale bar = 20 µm). The PCC was determined for the colocalization of the Alexa Fluor-568 (AGPAT1) and Alexa Fluor-488 (CHIKV E1) dyes. The right panel shows the line plot showing the colocalization of the green (CHIKV E1) and red (AGPAT1) fluorescence signals. (D) Huh7 cells were co-transfected with plasmids expressing AGPAT1-HA and CHIKV E2-FLAG. The cells were fixed, permeabilized and stained with rabbit anti-HA and mouse anti-FLAG antibodies. The cells were further incubated with anti-mouse Alexa Flour-488 and anti-rabbit Alexa-568 antibodies. The nuclei were stained with DAPI, and the cells were imaged using the Leica SP8 confocal microscope (scale bar = 20 µm). The PCC was determined for the colocalization of the Alexa Flour-488 (CHIKV E2) and Alexa Fluor-568 (AGPAT1) dyes. The right panel has the line plot for the two dyes. (E) Huh7 cells were transfected with the plasmids pAGPAT1, pE1-FLAG, or pcDNA5 as indicated. The cell lysates were prepared 48 h later and immunoprecipitated with anti-HA antibody. The pull-down and input samples were Western blotted with anti-HA antibody for detecting the AGPAT1-HA protein or anti-FLAG antibody for detecting the CHIKV E1-FLAG protein. (F) Huh7 cells were transfected with pAGPAT1-HA or pcDNA5, and 48 h later infected with CHIKV (MOI 3). The cell lysates were prepared 24 h pi and immunoprecipitated with anti-HA antibody. The pull-down and input samples were Western blotted with anti-HA antibody to detect the AGPAT1-HA protein and anti-E1 antibody to detect the CHIKV E1 protein. The representative microscopy images and Western blots are shown.

**Figure S6. The residue level interaction energy.**

The amino acid residues at the complex interface site (as numbered above) were analysed by the MM-GBSA approach. Per-residue energy breakdown analysis was used to determine the energy contribution of individual amino acids to identify critical residues at the interface and to reveal primary residue interactions within the complex by decomposing binding free energy (kcal/mol). Residues with higher interaction energies (cutoff =>3.5 kcal/mol) are identified; these are predicted to be involved in strong binding.

**Figure S7. Alanine scanning for the critical residue identification.**

The residue scanning was carried out by BioLuminate (Schrodinger) using the biologics residue scanning panel, side-chain prediction and backbone minimization as the refinement options, and a cutoff distance of 0.0. The net change in protein stability due to the mutation was calculated using the prime energy function, which included an implicit solvent term. The affinity change was calculated using Steepest (Schrodinger). The residues target for mutation to alanine (B and D) and the effect of the mutation on the complex affinity and stability is depicted in the plots (A and C). All energies are in kcal/mol.

**Figure S8. Role of AGPAT1 in CHIKV binding in ERMS cells.**

(A) ERMS cells were permeabilized with 0.3% Tween-20 or were not permeabilized. The cells were then incubated with rabbit AGPAT1 antibody and fixed with 2% paraformaldehyde followed by anti-rabbit Alexa Fluor-488 antibody. The nuclei were stained with DAPI, and the cells were imaged using the Leica SP8 confocal microscope (scale bar = 25 µm). The representative images are shown. (B) ERMS cells were incubated on ice with rabbit anti-AGPAT1 polyclonal antibody or rabbit IgG (isotype control) for 30 min at 10 µg/ml concentration, followed by incubation with CHIKV (MOI 50) on ice for 30 min. The cells were then incubated with CHIKV anti-E2 mouse monoclonal antibody on ice for 30 min and washed with ice-cold PBS. The cells were stained with Alexa Fluor-647 anti-mouse antibody for 30 min. The nuclei were stained with DAPI, and the cells were imaged using the Leica SP8 confocal microscope (scale bar = 25 µm). The right panel has the MIFD data. The representative images are shown. The statistical analysis of the data presented as mean±SD was done using the Student’s t-test with Welch’s correction; \*\*\**p*<0.001.

**Figure S9. Localization of AGPAT1 and MXRA8 on different cell lines.**

The cells were permeabilized with 0.3% Tween-20 or were not permeabilized, and incubated with polyclonal rabbit AGPAT1 or polyclonal rabbit MXRA8 antibody. The cells were then fixed, followed by incubation with anti-rabbit Alexa Fluor-568 antibody. The nuclei were stained with DAPI, and the cells were imaged using the Leica SP8 confocal microscope (scale bar = 25 µm, for zoom images scale bar = 10 µm). The representative images are shown.

## Acknowledgements

The work was supported by grant no. JCB/2021/000015 from the Science and Engineering Research Board, Govt. of India and benefitted from grant no. BT/MED/32/11/2019 from the Department of Biotechnology, Govt. of India to SV. BB received the Senior Research Fellowship (09/937(0014)/2019-EMR-I) from the Council of Scientific and Industrial Research (CSIR), India. SA and DS would like to acknowledge the NNP grant no. BT/PR40189/BTIS/137/50/2022 from the Department of Biotechnology, Govt. of India.

## Competing interest declaration

The authors declare no competing interests.

